# Sexual conflict promotes species coexistence through negative frequency dependence

**DOI:** 10.1101/2021.03.08.434376

**Authors:** Miguel Gomez-Llano, Sofie Nilén, Iain Moodie, Erik I. Svensson

## Abstract

A major challenge in community ecology is to understand the mechanisms promoting stable local coexistence. A necessary feature of local coexistence is that species show negative frequency dependence, rescuing rare species from exclusion. However, most studies have focused on ecological differences driving negative frequency dependence, ignoring non-ecological mechanisms such as reproductive interactions. Here, we combined field studies with behavioural and mesocosm experiments to investigate how reproductive interactions within and between species promote coexistence. Our results indicate that the intensity of male mating harassment and sexual conflict increases as species become more common, reducing female productivity and leading to negative frequency dependence. Moreover, field surveys reveal that negative frequency dependence operates in natural settings, consistent with our experimental results. These results suggest that sexual conflict can promote local coexistence and highlights the importance of studying reproductive interactions together with ecological differences to better understand the mechanisms promoting species coexistence.

**Significance statement:** Research on the mechanisms promoting local species coexistence have focused on canonical ecological differences that increase intraspecific over interspecific competition. However, one intrinsic factor of species that can promote coexistence are the reproductive interactions. We performed a series of behavioural and mesocosm experiments manipulating species frequencies together with field observations and show that sexual conflict can decrease female fitness when species are common and promote local coexistence. Our results suggest that reproductive interactions are an understudied mechanism that can promote species coexistence even when species are ecologically equivalent.

## Introduction

Understanding the causes underlying species diversity in ecological communities is a major challenge in both ecology and evolution. Coexistence theory predicts that negative frequency dependence is necessary for local species coexistence (1). If species have a fitness advantage when rare, they can increase from low abundance in a community and hence be rescued from competitive exclusion (1–3). Previous research has focused on how ecological differences between species can cause negative frequency dependence through rare species advantage (4–11), for example through predator susceptibility (8, 9), resource competition (5, 12) and phenology (11, 13). However, many communities are formed by species with little or no ecological differentiation (14–18). How or do such ecologically equivalent species coexist in a community? One possible answer to this question lays on an intrinsic characteristic of many species that can limit species population growth rate and promote species coexistence: reproductive interactions (19–22). Given how widespread sexual reproduction is in the tree of life, it is surprising how understudied reproductive interactions are as a mechanism for species coexistence.

Reproductive interactions can be categorized into four different groups: intraspecific interactions between the sexes, interspecific interactions between the sexes, intra- and interspecific interactions within the sexes. Importantly, not all reproductive interactions can promote local species coexistence. Intraspecific reproductive interactions between the sexes, such as male mating harassment and the resulting sexual conflict, can reduce female fitness and decrease population growth rate (23–26). Because male mating harassment and its fitness consequences for females is expected to be more intense when a species is common (23, 27–30), sexual conflict can in theory, promote local species coexistence (19, 20, 31, 32). Interspecific reproductive interactions between the sexes, when males mate or attempt to mate with females of another species (e. g., reproductive interference) (33) can lead to positive frequency dependence if females of the rare species suffer more from mating attempts by males of the common species, preventing local coexistence (34–36). Conversely, intraspecific competition within sexes (e. g., conspecific male-male competition) is expected to increase when a species is common (37). Moreover, male-male competition can affect male fitness by reducing longevity and/or male mating success (30, 38), leading to negative frequency dependence and local species coexistence (37, 39, 40). Finally, interspecific reproductive interactions within sexes (e.g., heterospecific male-male competition for mates) (41–43) can reduce male mating success and longevity (30, 37). Because the rare species will suffer more from heterospecific competition, such competition is expected to cause positive frequency dependence and prevent local species coexistence. Although male fitness is not always correlated with population growth, if males have reduced access to females, this could reduce the proportion of fertilized females and decrease population growth through reproductive collapse (44). Therefore, intra- and interspecific reproductive interactions within and between sexes can promote or prevent local species coexistence.

Importantly, these different types of reproductive interactions are likely to simultaneously operate within a given community. For example, males may compete both with conspecific and heterospecific males for mating territories (37). Because the frequency of these different reproductive interactions are likely to differ in importance and magnitude in different communities, studying only a subset of these interactions will only reveal a partial picture of community dynamics. For example, a previous study on *Calopteryx* damselflies suggested that in the presence of heterospecific male-male competition, the pressure from conspecific male mating harassment decreased and hence the intensity of sexual conflict (30). However, to the best of our knowledge no study has investigated all the different ways by which reproductive interactions within and between species can promote or prevent species coexistence.

Damselflies (Odonata: Zygoptera) have been widely used in research on sexual selection and sexual conflict, as they have intense reproductive interactions, such as conspecific male-male competition (37, 38, 45), heterospecific male-male competition (30, 37, 42, 43, 46), sexual conflict (25, 28, 30) and heterospecific matings and mating attempts (43, 47, 48). Moreover, they have been intensively used to investigate species coexistence in both the larvae (9, 10, 49–54) and less extensively adult stage (37). Therefore, these characteristics of damselflies make them ideal study systems to investigate the role of reproductive interactions in promoting local species coexistence.

Here, we used a combination of mating experiments, mesocosm experiments across the entire life cycle and surveys of natural damselfly assemblages to investigate if reproductive interactions can promote or prevent local species coexistence. Our focal study organisms are two species of pond damselflies, *Enallagma cyathigerum* and *Ischnura elegans* which are phenotypically very similar and they frequently co-occur (Fig. 1). Specifically we investigated 1) if any of the four types of reproductive interactions (intra- and interspecific within and between sexes) showed signs of frequency dependence; 2), if such reproductive interactions have a fitness cost; 3) if these two species are stably coexisting or only cooccurring (i.e., do they show negative frequency dependence); and 4) if these reproductive interactions and their fitness costs are likely to explain species frequencies and abundance dynamics across generations in natural communities. To answer these questions, we first carried out mating experiments where we manipulated species frequencies and test if the intensity of reproductive interactions changed when species are common compared to when rare. We proceed by quantifying the potential fitness costs of such reproductive interactions by measuring female survival and female productivity in a large multi-generational mesocosm with experimentally manipulated species frequencies. Finally, we quantified community dynamics through density- and frequency-changes at 18 communities across two generations. Taken together, our integrative study investigates if these species are stably coexisting, identifies sexual conflict as a mechanism that can promote local species coexistence, and shows that sexual conflict can explain species frequency changes across generations in natural settings. Our study therefore shows empirical evidence of a non-ecological mechanism promoting species coexistence, highlighting the need of broadening the views from traditional ecological perspectives and further integration of community ecology with evolutionary biology.

**Figure 1.**
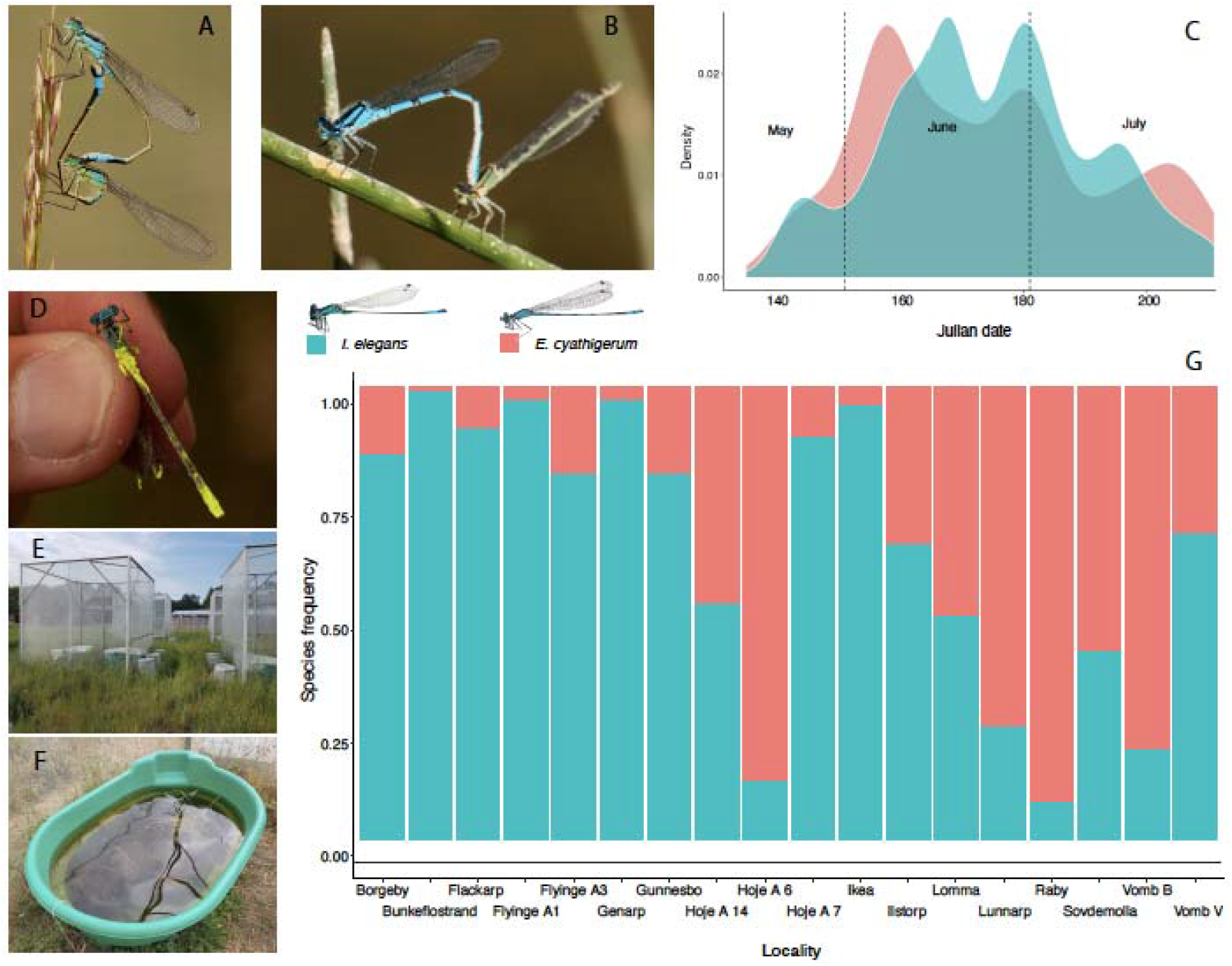
We used two species of damselflies *I. elegans* (**A**) and *E. cyathigerum* (**B**) to study the role of sexual conflict in species coexistence. These species are ecologically similar, and they overlap during the adult life stage (**C**). We performed mating experiments in small cages in which we marked male claspers and genitalia (**D**) to measure the intensity of different reproductive interactions when species are common and rare. We performed mesocosm experiments in large outdoor cages (**E**) with water containers (**F**) across the full life cycle to measure the costs of reproductive interactions. Finally, we did community surveys in 18 wild populations with different species frequencies (**G**) to estimate how species density changes were affected by initial species frequencies.

## Methods

### Study species

The common bluetail (*I. elegans;* Fig. 1A) and the common bluet (*E. cyathigerum;* Fig. 1B) are two ecologically similar and closely related damselfly species that are distributed throughout Eurasia with their northernmost range limits in Scandinavia, were they are commonly found in large numbers in ponds and lakes (55). These two damselfly species shared a most recent common ancestor at least 12.6 million years ago (56) and overlap extensively in their adult season (Fig. 1C) and are frequently locally sympatric (Fig. 1D). In southern Sweden where this study took place these species are univoltine (55), and due to their limited dispersal ability (less than 1Km) (57, 58), they can be found in a mosaic of largely discrete populations with different environmental conditions.

The reproductive behaviours of *Ischnura* and *Enallagma* are very similar. First, males are non-territorial and chase females (often several males at the time) and compete to grab the females by the prothorax using the claspers situated in the tip of their abdomens. If a male is able to find and subsequently clasp a female they form a tandem, after which the female can respond by bending the abdomen to reach the male genitalia and copulate (55) (Fig 1A). Before insemination the males remove the sperm from previous copulations (59, 60). Therefore, females gain few or no benefits from multiple matings, but will experience fitness costs that increase with the number of claspings (28). After copulating females oviposit in emergent vegetation and the larvae grow and overwinter in the aquatic stage (55). Males and females of both species are generalist predators and forage for flying insects near the water (55). During the reproductive season males interact frequently with both con- and heterospecific males, which can reduce male mating success (30, 37, 38, 42, 45, 48). In southern Sweden, where this study took place, adults of both species are found from late spring to late summer (late May to August) to reproduce (Fig 1 C).

### Are reproductive interactions frequency dependent?

To investigate if any of the four types of reproductive interactions shows frequency dependence, we carried out a mating trial experiment where we varied these two species relative frequencies: common (75%) and rare (25%). We used adult males and females (aged by the stiffness of the wings) from natural populations with no visible signs of external physical harm such as wing damage. We separated the captured individuals by sex and kept them at a density of 10 individuals in netted containers (10.2 cm diameter and 22.9 cm height) during transportation to Stensoffa Ecological Field Station, southern Sweden, where the experiments took place. At the field station we set up males and females in larger netted cages (45 cm diameter and 50 cm height). We added twigs and grasses to each cage to mimic natural vegetation and allow individuals to perch or rest, and a plastic cup with water to prevent desiccation. In each cage we put six individuals of one species (three males and three females) and two individuals of the other species (one male and one female). Thus, in these cages, we had two frequency treatments, both with equal sex ratios: “common” (75%) for the most abundant species and “rare” (25%) for the less abundant species (Supplementary Table 1A).

We marked all males in each cage with individual fluorescent colour powder in the genital area at the base of the abdomen and on the claspers (Fig 1B). After 24 hours, we terminated the experiment and searched for traces of colour dust on the genitalia and prothorax of the females. This technique allowed us to identify how many and which type of males (i.e., con-or heterospecifics) attempted to mate (i.e., clasped) or mated a given female, as these marked males left traces of colour dust in the female prothorax (i.e., mating attempt or clasping) and genitalia (i.e., successful mating) visible under UV-light. This method has previously been successfully used to quantify the degree of short-term male mating harassment (number of male claspings) and female mating rates in *I. elegans, E. cyathigerum* and other damselfly genera (28, 30, 37, 38).

To estimate the intensity of intraspecific reproductive interactions between sexes, and hence the potential for sexual conflict, we counted the number of male mating attempts (i.e., number of claspings per female) on conspecific females in 24 hours. We underscore that this rate of claspings does not take into account mating attempts that did not end up in claspings (i.e., chasing of females) or repeated claspings of females by the same male. Therefore, our measure of sexual conflict is conservative and will underestimate the total costs of male mating harassment to female fitness. To estimate interspecific reproductive interactions between sexes we counted the number of male mating attempts of heterospecific females, using the same procedure (i.e., remnants of coloured dust on the female prothorax or genitalia) in 24 hours. This measure is also a conservative measure of male mating harassment, as it does not take into account heterospecific mating attempts that did not end up in claspings. Finally, we quantified the costs of intra- and interspecific interactions within sexes as male mating success (mated = 1; not mated = 0). Because male-male competition can reduce male mating success (38), and if conspecific competition is stronger than heterospecific competition, male mating success is expected to decrease when species are common (i. e., negative frequency dependence). Conversely, if heterospecific male-male competition is stronger, male mating success is expected to decrease when species are rare (i. e., positive frequency dependence).

### Are reproductive interactions costly and do they cause negative frequency dependence and rare species fitness advantage?

We performed a series of mesocosm experiments under semi-natural conditions in eight large square outdoor cages (3m per side; total volume 27 m^3^) at the field station (Fig 1E). The aim of this mesocosm experiment was to quantify adult female longevity and per capita female productivity (i. e., the number of emerging female offspring in the next generation per female in the previous generation, a measure that should closely reflect population mean fitness or mean per capita growth rateunder different species frequency treatments (common, 75% and rare, 25%). Each cage contained a large water container (600L) with natural vegetation to resemble natural conditions and facilitate oviposition (Fig 1F). Each water container was inoculated repeatedly in the spring preceding these experiments with zooplankton (mainly copepods and *Daphnia)* obtained from nearby ponds and macrophytes obtained from an aquarium shop. This ensured that the damselfly larvae in our experiments would have enough food to forage and grow. A few weeks after inoculations we confirmed by visual inspections that these water containers had reproducing populations of zooplankton in the water. We added six coffee filter papers and small pieces of floating vegetation *(Phragmites australis)* to provide a resting substrate and to facilitate oviposition in these water containers. The outdoor cages were covered with mesh small enough to keep damselflies in and predators out, but wide enough to let smaller insects necessary as food for the foraging adults to enter (25, 30, 37). Importantly, these cages had no predators as we aimed to investigate if intra- and interspecific interactions could cause negative frequency dependence and potentially promote species coexistence. We have showed in previous studies that adult damselfly survival is not affected by total adult density, indicating that prey availability is not an issue in this experimental set up (30).

In each of these eight outdoor cages we manipulated species frequencies in two treatments with the same frequencies as in the mating trials described above: common (75%) and rare (25%). In each cage, we included 18 conspecifics (six females and 12 males) and six heterospecifics (two females and four males) for a total of 24 individuals per cage. We carried out a total of nine replicates (five for treatment with *I. elegans* being common and four for the treatment with *E. cyathigerum* being common) during the reproductive season (June and July) (Supplementary Table 2A). We also carried out two additional control treatments that would allow us to assess if there could be contamination in our water tanks from damselfly eggs attached to the vegetation or accidentally introduced through the zooplankton inoculation. The control cages contained 24 individuals (eight females and 16 males) of one species: two cages with only *I. elegans* (100 %) and two cages with only *E. cyathigerum* (100 %). Thus, any *E. cyathigerum* individuals that emerged in the *I. elegans* only control, and *vice versa* were considered as evidence for contamination from the outside. In each mesocosm cage, we marked every individual male and female with a unique number in two of the wings using permanent marker (such marking does not affect flight performance). This made possible measuring individual longevity by visiting these cages every day. We commonly observed marked females that mated and oviposited in the water containers inside the cages during the summer of 2018. In the two subsequent years (2019 and 2020), the cages were checked daily during the reproductive season to collect all emerging individuals, which were subsequently sexed and identified to species. Per capita female productivity per species was quantified as the number of female offspring emerged divided by the total number of adult females in the initial generation. Female productivity should be closely connected to female fitness and population growth rate (61).

### Do these species show negative frequency dependence in nature?

To test if our experiments show patterns consistent with assemblage dynamics (changes in species abundance) in natural populations, we surveyed 18 localities with natural *Ischnura-Enallagma* assemblages in southern Sweden during the reproductive season of 2018 and 2019, corresponding of two generations (Supplementary Table 3). These sites varied in relative species frequencies (Fig 1D). To quantify species frequencies and densities we visited each site between three and five times per season (May-July) during warm (>15°C) days with no rain or strong wind, the most favourable conditions for these damselflies (Supplementary Table 3). During these visits, we captured as many individuals as possible with hand nets for 30 minutes, after which each individual was sexed and identified to species. The relative frequency of each species was taken as the number of individuals of that species divided by the total number of individuals of both species in each season. Species densities were calculated as encountering rate, number of individuals caught per sampling time (i. e., individuals caught per person-minute). Encountering rate has been used previously as a proxy of species density in adult damselflies (28). We calculated the changes in species densities across years (generations) by dividing the density of each species at a given site in 2019 to the initial density of the same species in 2018 (number of adults in 2019 per adult in 2018).

### Statistical analysis

Statistical analyses were carried out using generalized linear models assuming poisson (number of con- and heterospecific claspings), binomial (male mating success), negative binomial (longevity) and normal (female productivity) distributions of the residuals. The number of con- and heterospecific claspings, male mating success, female longevity and productivity were all treated as dependent variables. Species frequency, species and their interaction were included as fixed factors. In the analysis of number of con- and heterospecific claspings and male mating success in the mating experiments, we controlled for replicate cage number including it as a covariate. In the model of adult female longevity in the mesocosm experiments, we included the interaction between cage and replicate as random factors. Species density changes in the 18 natural communities was analysed using a linear model with initial species frequencies in 2018, species and their interaction as fixed factors. All models were performed using the packages “lme4” (62) and “car” (63) in R (64).

## Results

### Are reproductive interactions frequency dependent?

We quantified the number of conspecific and heterospecific claspings from 87 females and mating success of 89 males in our mating trials. We found a significant effect of species frequency on the number of con- and heterospecific claspings of females, but in opposite directions. Females experienced more conspecific claspings when they were common than when they were rare (χ^2^ = 4.61, *p* = 0.031; Fig. 2A) but more heterospecific claspings when they were rare than when they were common (χ^2^ = 11.12, *p* < 0.001; Fig. 2B). In contrast, we found no effect of species frequency on male mating success. In all the models we found no effect of species nor the interaction between species and frequency (Supplementary Table 1B-D).

**Figure 2.**
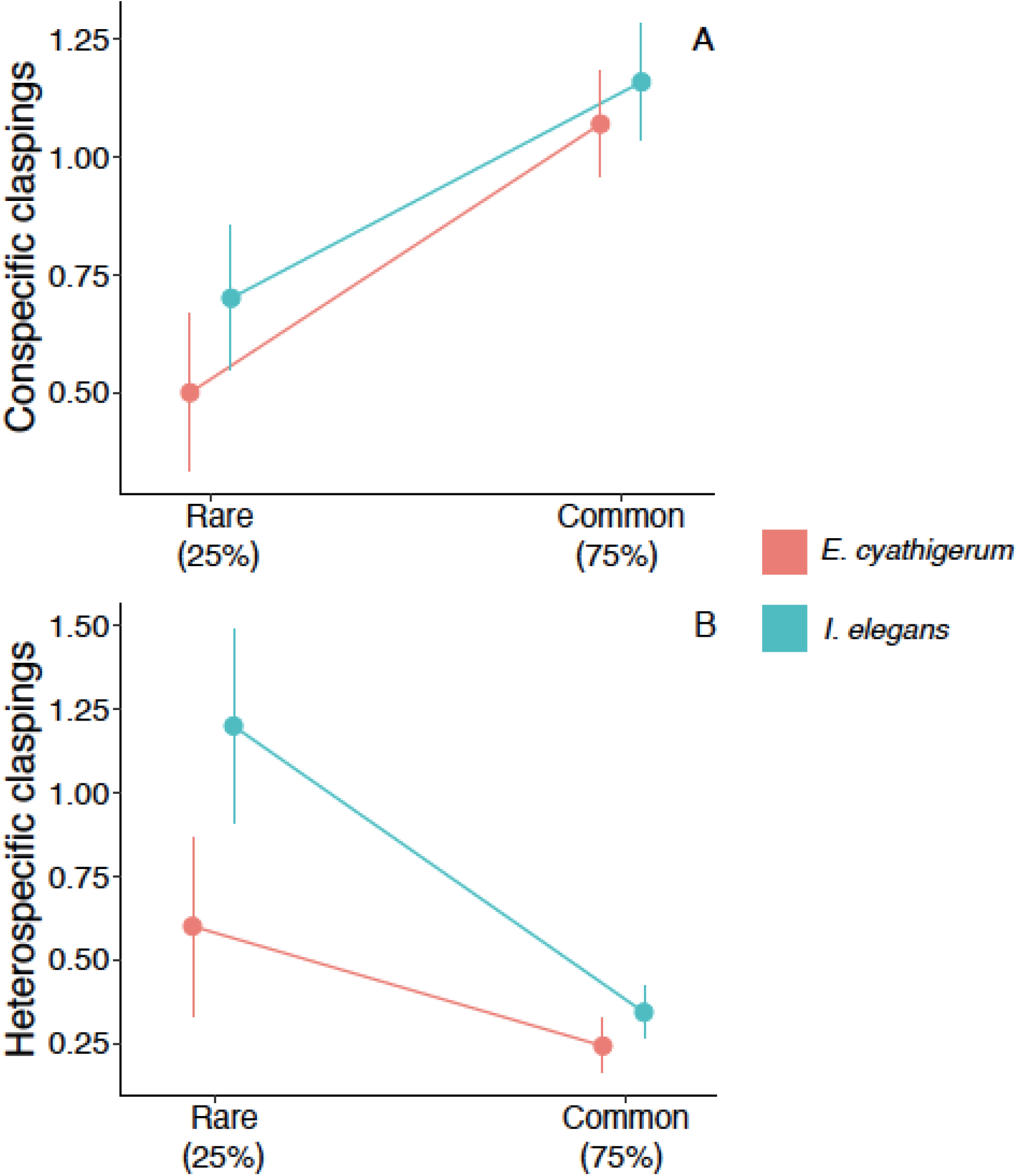
We perform a series of mating experiments in which we manipulated species frequencies, “rare” (25%) and “common (75%) to test the intensity of intra and interspecific reproductive interactions (Supplementary Table 1). We found that the intensity of sexual conflict, measured as the number of mating attempts (i.e., claspings), was more intense when species were common than when rare (**A**). Heterospecific matings attempts followed the opposite pattern, females experiencing more mating attempts by heterospecifics when rare than when common (**B**). Points show the means and error bars the standard error.

### Are reproductive interactions costly and do they result in rare species advantage?

We quantified adult longevity for 128 females (64 of each species) in our mesocosm experiments. We found no main effect of frequency treatment on female longevity nor a significant interaction between species and frequency, suggesting that neither species longevity was affected by changes in the species frequency. However, we found a significant main effect of species on female longevity (χ^2^ = 24.72, *p* < 0.001), with shorter longevity (> 50%) of *E. cyathigerum* compared to *I. elegans* in these mesocosm cages (Supplementary Table 2B).

Next, we analysed female productivity (i.e., the number of female offspring in the next generation per adult female in the initial generation) in the mesocosm experiments. Female productivity differed significantly between the two species (F = 15.63, *p* = 0.028), with *I. elegans* females being on average more productive than *E. cyathigerum.* Importantly, we found a significant and negative effect of species frequency (F = 53.55, *p* = 0.005) on female productivity, with lower female productivity in the common compared to the rare frequency treatment (i.e., negative frequency dependence) (Fig. 3). We found no significant interaction between species identity and species frequency (Supplementary Table 2C), suggesting that the strength of negative frequency-dependence was similar in both species. The results were similar when we analysed the total number of emerging individuals in the offspring generation and the number of emerging male offspring (Supporting Table 2D). We found only minor contamination in our control cages, and our results above remain qualitatively similar after correcting for such contamination (Supplementary Analysis 1).

**Figure 3.**
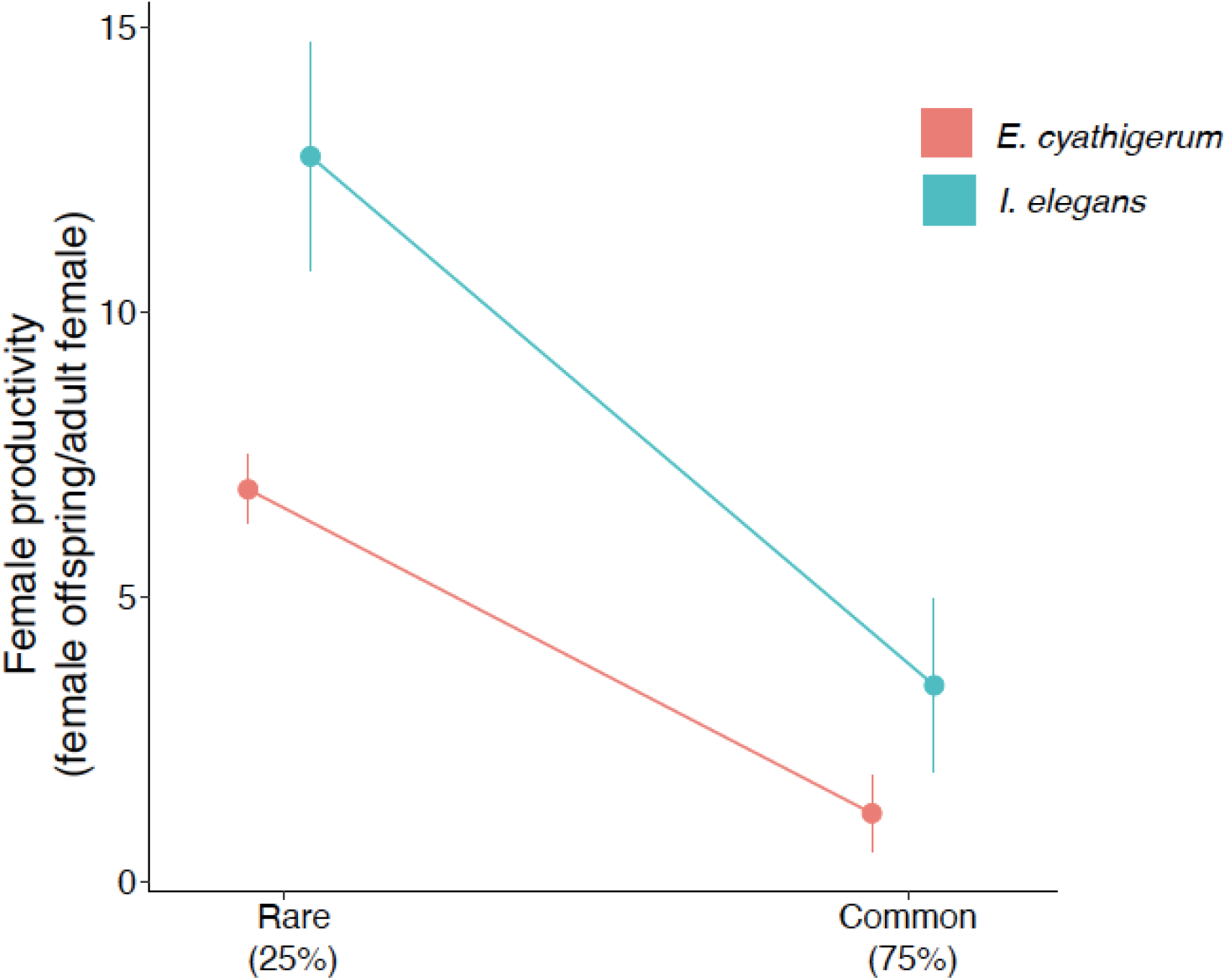
We used mesocosm experiments to quantify the costs of sexual conflict in female fitness. We found strong negative frequency-dependence in female productivity (measured as the number of female offspring that emerge in the following generation per female in the initial generation), having higher fitness when rare over common. Similar results were found when we analyzed total productivity (i.e., number of offspring emerged per adult female in the initial generation; Supplementary Table 2). Points show the means and error bars the standard error.

### Do these species show negative frequency dependence in nature?

Finally, we analysed species density changes across two generations at the18 natural sympatric sites of *I. elegans* and *E. cyathigerum.* We found a significant effect of initial species frequency (we present results on a logarithmic scale as they show better fit, although untransformed data was also significant) on species density change (F = 14.95, *p* < 0.001; Fig. 4). There was no significant effect of species identity nor the interaction between species identity and initial frequency (Supplementary Table 4), suggesting that these two species respond similarly to changes in relative frequencies in nature. These between-generation changes indicate that the higher species frequency was at a site in 2018, the more it declined in abundance the following year. These results suggest negative frequency dependence, consistent with the findings in the mesocosm experiment (Fig. 3).

**Figure 4.**
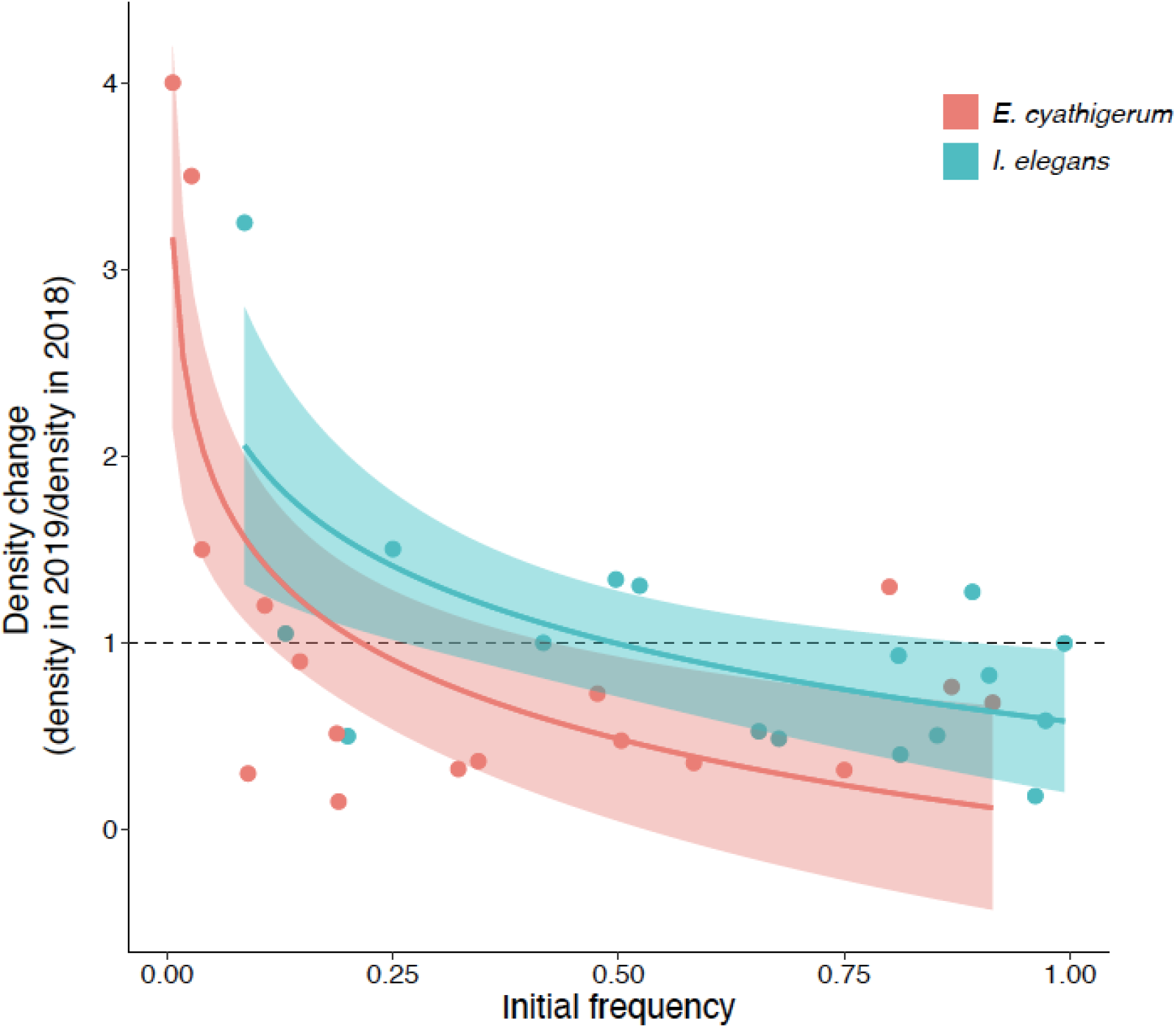
We monitored damselfly communities in 18 localities during two consecutive years (2018-2019; Supplementary Table 3). We calculated the initial species frequency in 2018 and the species density change (i.e., species density in 2019/ density in 2018). We found evidence for negative frequency dependence and rare species advantage in both *I. elegans* and *E. cyathigerum* (Supplementary Table 4). Higher the initial species frequency in 2018 suffer a density decline, lower initial frequencies increase their densities. Points show individuals observations at each locality, line show model predictions (best fit with in a logarithmic regression) and shaded areas confidence interval.

## Discussion

Negative frequency dependence is an fundamental requirement for species coexistence, as a species that has a fitness advantage when rare can recover from low abundance and competitive exclusion can be prevented (1, 2). However, our understanding of the underlying mechanisms responsible for negative frequency dependence and stable coexistence is still poor (2, 65). Many previous studies have focused on the ecological niche differences by which negative interspecific interactions can be reduced, although reproductive interactions alone can also cause negative frequency dependence and promote species coexistence (19–22, 32). Our results suggest that intraspecific male mating harassment and the resulting sexual conflict it generates has the potential to reduce female productivity, causing negative frequency dependence and promoting species coexistence. To the best of our knowledge, our study is the first empirical example of how a mechanism not based on ecological niche differences can promote stable species coexistence.

Sexual conflict can have severe negative effects on female fitness and by extension reduce population growth (23–27, 66–70). Moreover, sexual conflict is expected to increase at higher densities (25, 27, 28, 30), as high densities should increase encounter rates between the sexes and thereby elevate male mating harassment on females (25, 28). If the negative fitness effects of sexual conflict on females are larger when species are common and reduced when rare, sexual conflict could lead to negative frequency-dependence and rescue rare species from competitive exclusion (19, 20). Consistent with these predictions, we found evidence for negative frequency-dependence on female productivity (Fig. 3). Our mating experiments suggest that sexual conflict could be the driving mechanism causing negative frequency dependence. Although other mechanisms (e.g., predation, cannibalism), especially during larvae stage could also influence species relative frequency changes in the wild. Conversely, heterospecific mating attempts are likely to be shorter in duration, given that females reject heterospecific males and given that heterospecific male claspers do typically not match female prothorax structures (71, 72). Therefore, heterospecific mating attempts are likely to be less costly than conspecific male mating attempts.

We suggest that the loss of female productivity in our mesocosm experiment corresponds to a doubling of male mating harassment, measured as the number of conspecific claspings in both *E. cyathigerum* and *I. elegans* when they are common (Fig. 2A). In our mesocosm experiments, female productivity of *E. cyathigerum* when common was only 17% of the productivity when rare, and in *I. elegans* productivity when common was only 27% of the productivity when rare, demonstrating strong negative frequency dependence of female fitness. Moreover, in damselflies, males remove the sperm from previous copulations, leading to no obvious female benefits from multiple matings (59, 60). Our experimental setup only allowed us to identify if a male attempted to mate a female if he managed to clasp her, but we could not quantify mating attempts that did not result in claspings (i.e., chasing and fighting), nor if there were multiple mating attempts by the same male. Moreover, the effect of male harassment on females might not translate only in loss of fecundity but also reduce larvae survival (73). Therefore, our measure of sexual conflict is likely to be conservative and underestimate the true level of total mating harassment that females are likely to have experienced in our experimental settings and in nature.

In addition to male mating harassment and sexual conflict in the adult stage, intra- and interspecific competition during the aquatic larval stage could also potentially have decreased the number of adult female emergences. However, a previous study found that under the current environmental conditions, *I. elegans* and *E. cyathigerum* larvae were competitive equivalent (54). Moreover, previous experimental evidence with different *Ischnura* and *Enallagma* species from North America showed no frequency-dependent mortality or growth rate in larvae in the absence of predators (52). In agreement with our mesocosms with no predators *(but see* (50) for an effect of relative frequency in growth rates). However, in natural settings damselfly species coexistence can be achieved by predation during larvae causing negative frequency dependence (9, 52). Moreover, *Ischnura* and *Enallagma* species in North America show striking larval behavioural differences, with *Ischnura* being more active and susceptible to fish or dragonfly predators (53). Given that damselflies in our area occur in a mosaic of lakes with fish or dragonflies as the top predator, as well as in the absence of them, it is likely that different mechanisms could be acting separately or synergistically in the different localities. However, in the absence of top predators, sexual conflict could have a sufficiently large effect to rescue populations from exclusion and promote coexistence.

We have experimentally investigated how sexual conflict can affect species coexistence and by extension the maintenance of local diversity. Our results point to the importance of mechanisms based on the intrinsic reproductive interactions within species and suggest that sexual conflict can generate negative frequency dependence and promote species coexistence. However, further evaluating the relative importance and interaction of reproductive interactions with other ecological factors, such as the presence and type of predators, would help us better understand the mechanisms promoting species coexistence in nature.

## Acknowledgements

Adam Siepielski for critical comments on earlier versions of this manuscript. MGL was supported by NSF DEB1748945 and The Royal Physiographic Society of Lund. ES was supported by The Swedish Research Council (2016-03356), Carl Tryggers Fundtion, Gyllenstiernska Krapperupstiftelsen (KR2018-0038), Lunds Djurskyddsfond, The Royal Physiographic Society of Lund, Stiftelsen Olle Engqvist Byggmästare, and the Stina Werners Foundation. SN was supported by the Sven and Lili Lawski’s Foundation.

## Authors contributions

MGL and ES design the study. All authors carried out the experiments. MGL perform the analysis and wrote the text with input from all the authors.

## Supplementary Information

**Supplementary Table 1.** We performed a series of behavioural mating assays in which we quantified con- and heterospecific mating attempts and male mating success under different frequency treatments. The sample sizes in the different frequency treatments, number of individuals used of each sex and number of replicates are shown in (**A**). We analysed the effects of species frequency, species and their interaction in conspecific claspings (**B**), heterospecific claspings (**C**) and for male mating success (**D**).

| Species frequency | Species | No. males | No. females | Replicates | Tot. males | Tot. females |
| --- | --- | --- | --- | --- | --- | --- |
| Common (75%) | *I. elegans* | 3 | 3 | 13 | 39 | 39 |
| Rare (25%) | *E. cyathigerum* | 1 | 1 |  | 13 | 13 |
| Common (75%) | *E. cyathigerum* | 3 | 3 | 10 | 30 | 30 |
| Rare (25%) | *I. elegans* | 1 | 1 |  | 10 | 10 |

B) Number of conspecific male claspings.
| | $\chi^2$ | df | P |
| --- | --- | --- | --- |
| Replicate | 0.25 | 1 | 0.614 |
| Frequency | 4.61 | 1 | **0.031** |
| Species | 0.44 | 1 | 0.504 |
| Frequency : Species | 0.08 | 1 | 0.772 |

C) Number of heterospecific claspings.
| | $\chi^2$ | df | P |
| --- | --- | --- | --- |
| Replicate | 0.007 | 1 | 0.933 |
| Frequency | 11.12 | 1 | **<0.001** |
| Species | 2.38 | 1 | 0.122 |
| Frequency : Species | 0.207 | 1 | 0.648 |

D) Male mating success (mated=1; un-mated=0).
| | $\chi^2$ | df | P |
| --- | --- | --- | --- |
| Replicate | 0.34 | 1 | 0.559 |
| Frequency | 0.271 | 1 | 0.602 |
| Species | 0.149 | 1 | 0.699 |
| Frequency : Species | 0.148 | 1 | 0.699 |

**Supplementary Table 2.** We carried out multi-generational mesocosm experiments in large outdoor cages with water containers across the entire life-cycle of damselflies (Fig. 1). We estimated female productivity (number of emerging female offspring in the next generation per adult female in the initial generation) for the two different species *(I. elegans* and *E. cyathigerum)* under the two different frequency treatments (Rare: 25 % and Common: 75 %). Sample sizes in the different frequency treatments, number of individuals per replicate and in total and the number of replicates are shown in (**A**). We analysed the effect of species frequency, species and their interaction on female longevity (**B**), female productivity (measured as the number of female offspring emerged in the next generation per adult female in the parental generation in 2018; **C**), and total female productivity (measured as the total number of offspring emerged in the next generation per adult female in the parental generation; **D**).

| Species frequency | Species | No. males | No. females | Replicates | Tot. males | Tot. females |
| --- | --- | --- | --- | --- | --- | --- |
| Common (75%) | *I. elegans* | 12 | 6 | 5 | 60 | 30 |
| Rare (25%) | *E. cyathigerum* | 4 | 2 |  | 20 | 10 |
| Common (75%) | *E. cyathigerum* | 12 | 6 | 4 | 48 | 24 |
| Rare (25%) | *I. elegans* | 4 | 2 |  | 16 | 8 |

B) Adult female longevity (days).
| | $\chi^2$ | df | P |
| --- | --- | --- | --- |
| Frequency | 0.116 | 1 | 0.733 |
| Species | 24.724 | 1 | < **0.001** |
| Frequency : Species | 0.1003 | 1 | 0.751 |

C) Female per-capita productivity (No. emerging female offspring in 2019 and 2020).
|  | F | df | P |
| --- | --- | --- | --- |
| Frequency | 31.68 | 1 | **0.004** |
| Species | 9.25 | 1 | **0.038** |
| Frequency : Species | 1.83 | 1 | 0.247 |

|  | F | df | P |
| --- | --- | --- | --- |
| Frequency | 18.29 | 1 | **0.012** |
| Species | 5.11 | 1 | 0.086 |
| Frequency : Species | 1.2 | 1 | 0.333 |

**Supplementary Table 3.**
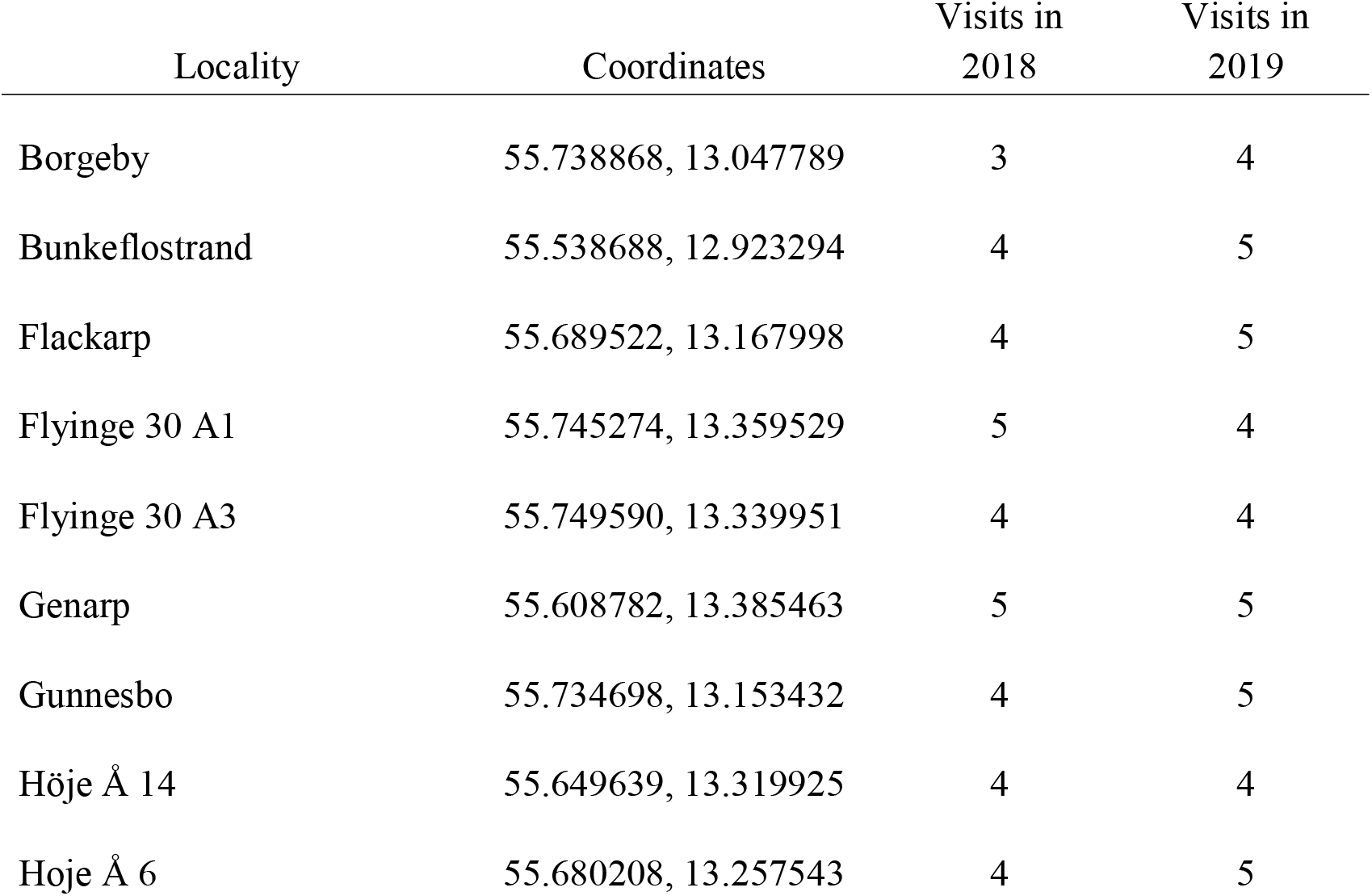
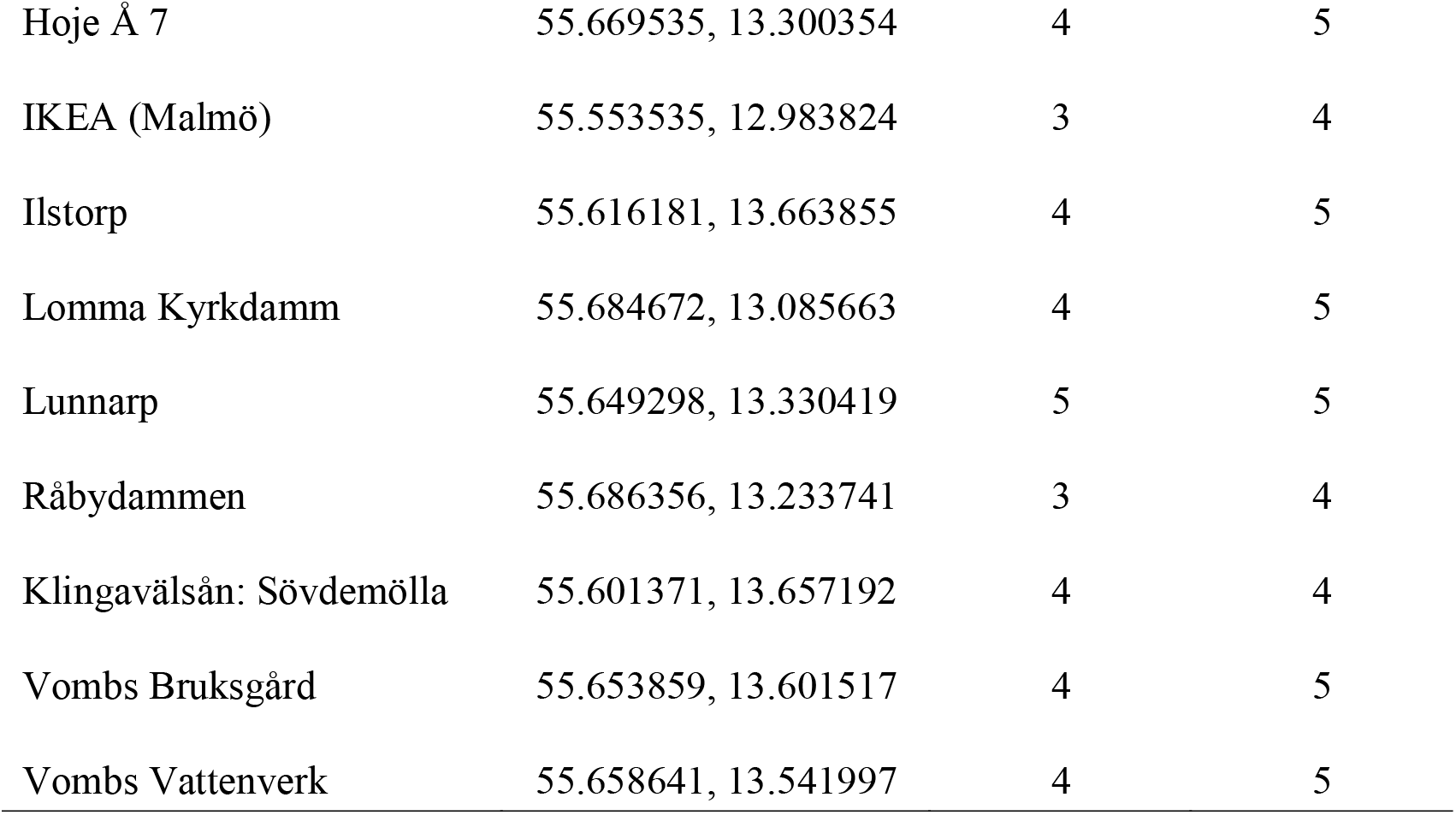
We monitored community dynamics at 18 natural sites during the reproductive season of 2018 and 2019, quantifying the relative species frequency and density in both years. Here we provide information about the geographic locations of these different sites.

| Locality | Coordinates | Visits in<br>2018 | Visits in<br>2019 |
| --- | --- | --- | --- |
| Borgeby | 55.738868, 13.047789 | 3 | 4 |
| Bunkeflostrand | 55.538688, 12.923294 | 4 | 5 |
| Flackarp | 55.689522, 13.167998 | 4 | 5 |
| Flyinge 30 A1 | 55.745274, 13.359529 | 5 | 4 |
| Flyinge 30 A3 | 55.749590, 13.339951 | 4 | 4 |
| Genarp | 55.608782, 13.385463 | 5 | 5 |
| Gunnesbo | 55.734698, 13.153432 | 4 | 5 |
| Höje Å 14 | 55.649639, 13.319925 | 4 | 4 |
| Hoje Å 6 | 55.680208, 13.257543 | 4 | 5 |
| Hoje Å 7 | 55.669535, 13.300354 | 4 | 5 |
| IKEA (Malmö) | 55.553535, 12.983824 | 3 | 4 |
| Ilstorp | 55.616181, 13.663855 | 4 | 5 |
| Lomma Kyrkdamm | 55.684672, 13.085663 | 4 | 5 |
| Lunnarp | 55.649298, 13.330419 | 5 | 5 |
| Råbydammen | 55.686356, 13.233741 | 3 | 4 |
| Klingavälsån: Sövdemölla | 55.601371, 13.657192 | 4 | 4 |
| Vombs Bruksgård | 55.653859, 13.601517 | 4 | 5 |
| Vombs Vattenverk | 55.658641, 13.541997 | 4 | 5 |

**Supplementary Table 4.** Community dynamics in the 18 wild populations was analysed as the effect of species frequency in 2018 (log-transformed; **A**) and species identity in species density change (density in 2019/density in 2018) using a general linear model. Nontransformed species frequency in 2018 show qualitatively similar results (**B**).

**A**
|  | F | df | p |
| --- | --- | --- | --- |
| log(Frequency 2018) | 14.95 | 1 | < **0.001** |
| Species | 0.438 | 1 | 0.512 |
| log(Frequency 2018) : Species | 0.932 | 1 | 0.341 |
| Adj. R<sup>2</sup> = 0.286 |  |  |  |

**B**
|  | F | df | p |
| --- | --- | --- | --- |
| Frequency 2018 | 18.39 | 1 | **0.041** |
| Species | 0.005 | 1 | 0.972 |
| Frequency 2018 : Species | 5.56 | 1 | 0.25 |
| Adj. R<sup>2</sup> = 0.098 |  |  |  |

**Supplementary Analysis 1.** We found evidence of small contamination of both species in the control cages with allopatric species frequencies (100 % *I. elegans* or *E. cyathigerum).* The source of this contamination might have come from larvae or eggs attached to the vegetation used to inoculate the water containers or adult of either species entering the cages by mistake. The mean contamination in the two *E. cyathigerum* cages was 18% (min = 5%, max = 31%), and in the two *I. elegans* cages 2% (min = 0.8%, max = 4%). Assuming similar levels of contamination occurred in the other cages, we performed two corrections to see how this might have confounded our results. First, we removed from the female productivity of each cage the mean percentage of contamination of each species, (−18% and −2% of *E. cyathigerum* and *I. elegans,* respectively) (**A**). Second, we performed a more conservative correction, by instead removing the maximum level of contamination found of each species (31% and −4% of *E. cyathigerum* and *I. elegans,* respectively), (**B**). The two models below (**A** and **B**) were performed using the corrected value of female productivity as response variable, species frequency, species identity and their interaction as fixed factors in a generalized linear model. In both cases the results are qualitatively the same as in the uncorrected emergences and the effects of both “frequency” and “species” remain highly significant (see also Fig. 3).

|  | F | df | p |
| --- | --- | --- | --- |
| Frequency | 28.91 | 1 | **0.005** |
| Species | 12.88 | 1 | **0.022** |
| Frequency : Species | 2.97 | 1 | 0.159 |

B) Corrected analysis of female productivity assuming maximum contamination equivalent to the control cages.
|  | F | df | p |
| --- | --- | --- | --- |
| Frequency | 26.72 | 1 | **0.006** |
| Species | 16.1 | 1 | **0.015** |
| Frequency : Species | 4.05 | 1 | 0.114 |

## References

1. P. Chesson, Mechanisms of maintenance of species diversity. Annu. Rev. Ecol. Syst. 31, 343–366 (2000).

2. P. B. Adler, J. HilleRisLambers, J. M. Levine, A niche for neutrality. Ecol Lett 10, 95–104 (2007).

3. R. MacArthur, R. Levins, The limiting similarity, convergence, and divergence of coexisting species. Am. Nat. 101, 377–385 (1967).

4. F. J. Ayala, Competition between species: frequency dependence. Science 171, 820–824 (1971).

5. R. H. MacArthur, Geographical ecology: patterns in the distribution of species (Princeton University Press, 1972).

6. S. J. Hausch, J. W. Fox, S. M. Vamosi, Coevolution of competing *Callosobruchus* species does not stabilize coexistence. Ecol. Evol. 7, 6540–6548 (2017).

7. C. Wills, et al., Nonrandom processes maintain diversity in tropical forests. Science 311, 527–531 (2006).

8. P. Chesson, J. J. Kuang, The interaction between predation and competition. Nature 456, 235–238 (2008).

9. J. T. Bried, A. M. Siepielski, Predator driven niches vary spatially among co-occurring damselfly species. Evol. Ecol. 33, 243–256 (2019).

10. B. H. Ousterhout, M. Serrano, J. T. Bried, A. M. Siepielski, A framework for linking competitor ecological differences to coexistence. J. Anim. Ecol. (2019).

11. C. Blackford, R. M. Germain, B. Gilbert, Species differences in phenology shape coexistence. Am. Nat. 195, E168–E180 (2020).

12. T. W. Schoener, Resource partitioning in ecological communities. Science 185, 27–39 (1974).

13. J. Usinowicz, et al., Temporal coexistence mechanisms contribute to the latitudinal gradient in forest diversity. Nature 550, 105–108 (2017).

14. C. Boake, Sexual selection and speciation in Hawaiian *Drosophila*. Behav. Genet. 35 (2005).

15. W. Salzburger, The interaction of sexually and naturally selected traits in the adaptive radiations of cichlid fishes. Molec. Ecol. 18, 169–185 (2009).

16. T. Price, I. J. Lovette, E. Bermingham, H. L. Gibbs, A. D. Richman, The Imprint of History on Communities of North American and Asian Warblers. Am. Nat. 156, 354–367 (2000).

17. M. A. McPeek, J. M. Brown, Building a regional species pool: diversification of the Enallagma damselflies in eastern North America. Ecology 81, 904 (2000).

18. A. M. Siepielski, A. Z. Hasik, B. H. Ousterhout, An ecological and evolutionary perspective on species coexistence under global change. Curr. Opin. Ins. Scien. 29, 71–77 (2018).

19. D.-Y. Zhang, I. Hanski, Sexual reproduction and stable coexistence of identical competitors. J. Theor. Biol. 193, 465–473 (1998).

20. K. Kobayashi, Sexual selection sustains biodiversity via producing negative density dependent population growth. J. Ecol. 107, 1433–1438 (2019).

21. OsamuK. Mikami, M. Kohda, M. Kawata, A new hypothesis for species coexistence: male-male repulsion promotes coexistence of competing species. Popul Ecol 46 (2004).

22. L. Ruokolainen, I. Hanski, Stable coexistence of ecologically identical species: conspecific aggregation via reproductive interference. J Anim Ecol 85, 638–647 (2016).

23. H. Kokko, R. Brooks, Sexy to die for? Sexual selection and the risk of extinction in Annales Zoologici Fennici, (JSTOR, 2003), pp. 207–219.

24. J.-F. Le Galliard, P. S. Fitze, R. Ferrière, J. Clobert, Sex ratio bias, male aggression, and population collapse in lizards. PNAS 102, 18231–18236 (2005).

25. Y. Takahashi, K. Kagawa, E. I. Svensson, M. Kawata, Evolution of increased phenotypic diversity enhances population performance by reducing sexual harassment in damselflies. Nat Commun 5, 4468 (2014).

26. L. Yun, et al., Competition for mates and the improvement of nonsexual fitness. PNAS 115, 6762–6767 (2018).

27. H. Kokko, D. J. Rankin, Lonely hearts or sex in the city? Density-dependent effects in mating systems. Phil. Trans. R Soc B 361, 319–334 (2006).

28. T. P. Gosden, E. I. Svensson, Density-dependent male mating harassment, female resistance, and male mimicry. Am Nat 173, 709–721 (2009).

29. O. Y. Martin, D. J. Hosken, The evolution of reproductive isolation through sexual conflict. Nature 423, 979–982 (2003).

30. M. A. Gomez-Llano, H. M. Bensch, E. I. Svensson, Sexual conflict and ecology: Species composition and male density interact to reduce male mating harassment and increase female survival. Evolution 72, 906–915 (2018).

31. K. Kobayashi, Sexual reproduction and diversity: Connection between sexual selection and biological communities via population dynamics. Popul Ecol 61, 135–140 (2019).

32. M. Yamamichi, et al., Intraspecific adaptation load: a mechanism for species coexistence. Trend Ecol Evol (2020).

33. G. F. Grether, K. S. Peiman, J. A. Tobias, B. W. Robinson, Causes and consequences of behavioral interference between species. Trend Ecol Evol 32, 760–772 (2017).

34. E. Kuno, Competitive exclusion through reproductive interference. Res Popul Ecol 34, 275–284 (1992).

35. P. Amarasekare, Interference competition and species coexistence. Proc R Soc Lond B 269, 2541–2550 (2002).

36. J. Whitton, C. J. Sears, W. P. Maddison, Co-occurrence of related asexual, but not sexual, lineages suggests that reproductive interference limits coexistence. Proc. R. Soc. B 284, 20171579 (2017).

37. E. I. Svensson, M. A. Gómez-Llano, A. R. Torres, H. M. Bensch, Frequency dependence and ecological drift shape coexistence of species with similar niches. Am Nat 191, 691–703 (2018).

38. M. Gómez-Llano, A. Narasimhan, E. I. Svensson, Male-male competition causes parasite-mediated sexual selection for local adaptation. Am Nat 196, 344–354 (2020).

39. M. Kohda, Does male-mating attack in the herbivorous cichlid, *Petrochromis polyodon,* facilitate the coexistence of congeners? Ecol Freshw Fish 4, 152–159 (1995).

40. M. Kohda, “Coexistence of permanently territorial cichlids of the genus *Petrochromis* through male-mating attack” in Fish Biology in Japan: An Anthology in Honour of Hiroya Kawanabe, (Springer, 1998), pp. 231–242.

41. K. A. Dixon, W. H. Cade, Some factors influencing male-male aggression in the field cricket *Gryllus integer* (time of day, age, weight and sexual maturity). Anim Behav 34, 340–346 (1986).

42. J. P. Drury, G. F. Grether, Interspecific aggression, not interspecific mating, drives character displacement in the wing coloration of male rubyspot damselflies *(Hetaerina)*. Proc. R Soc. B 281, 20141737 (2014).

43. J. P. Drury, et al., A general explanation for the persistence of reproductive interference. Am Nat 194, 268–275 (2019).

44. E. J. Milner-Gulland, et al., Reproductive collapse in saiga antelope harems. Nature 422, 135–135 (2003).

45. G. F. Grether, J. P. Drury, E. Berlin, C. N. Anderson, The role of wing coloration in sex recognition and competitor recognition in rubyspot damselflies *(Hetaerina* spp.). Ethology 121, 674–685 (2015).

46. K. Tynkkynen, M. J. Rantala, J. Suhonen, Interspecific aggression and character displacement in the damselfly *Calopteryx* splendens. J Evol Biol 17, 759–767 (2004).

47. E. I. Svensson, K. Karlsson, M. Friberg, F. Eroukhmanoff, Gender differences in species recognition and the evolution of asymmetric sexual isolation. Curr Biol 17, 1943–1947 (2007).

48. J. P. Drury, K. W. Okamoto, C. N. Anderson, G. F. Grether, Reproductive interference explains persistence of aggression between species. Proc. R. Soc. B 282, 20142256–20142256 (2015).

49. A. M. Siepielski, K.-L. Hung, E. E. B. Bein, M. A. McPeek, Experimental evidence for neutral community dynamics governing an insect assemblage. Ecology 91, 847–857 (2010).

50. A. M. Siepielski, A. N. Mertens, B. L. Wilkinson, M. A. McPeek, Signature of ecological partitioning in the maintenance of damselfly diversity. J Anim Ecol 80, 1163–1173 (2011).

51. A. M. Siepielski, A. Nemirov, M. Cattivera, A. Nickerson, Experimental evidence for an eco-evolutionary coupling between local adaptation and intraspecific competition. Am Nat 187, 447–456 (2016).

52. M. A. McPeek, The consequences of changing the top predator in a food web: a comparative experimental approach. Ecol Monog 68, 1–23 (1998).

53. M. A. McPeek, The growth/predation risk trade-off: so what is the mechanism? Am Nat 163, E88–E111 (2004).

54. L. T. Lancaster, G. Morrison, R. N. Fitt, Life history trade-offs, the intensity of competition, and coexistence in novel and evolving communities under climate change. Phil Trans R Soc B 372, 20160046 (2017).

55. P. S. Corbet, Dragonflies: behaviour and ecology of Odonata (Harley Books, 1999).

56. J. Swaegers, et al., Ecological and evolutionary drivers of range size in *Coenagrion* damselflies. J Evol Biol 27, 2386–2395 (2014).

57. K. F. Conrad, et al., Characteristics of dispersing Ischnura elegans and *Coenagrion puella* (Odonata): age, sex, size, morph and ectoparasitism. Ecography 25, 439–445 (2002).

58. K. F. Conrad, K. H. Willson, I. F. Harvey, C. J. Thomas, T. N. Sherratt, Dispersal characteristics of seven odonate species in an agricultural landscape. Ecography 22, 524–531 (1999).

59. O. M. Fincke, Sperm competition in the damselfly *Enallagma hageni* Walsh (Odonata: Coenagrionidae): benefits of multiple mating to males and females. Behav Ecol Sociobiol 14, 235–240 (1984).

60. A. Córdoba-Aguilar, E. Uhía, A. C. Rivera, Sperm competition in Odonata (Insecta): the evolution of female sperm storage and rivals’ sperm displacement. J. Zoology 261, 381–398 (2003).

61. H. Caswell, Analysis of life table response experiments I. Decomposition of effects on population growth rate. Ecological Modelling 46, 221–237 (1989).

62. D. Bates, M. Mächler, B. Bolker, S. Walker, Fitting linear mixed-effects models using lme4. J Statis Software 67, 1–48 (2015).

63. J. Fox, S. Weisberg, An {R} companion to applied regression, Third (Sage, 2011).

64. R Development Core Team, R: A language and environment for statistical computing (R foundation for statistical computing Vienna, Austria, 2018).

65. M. A. McPeek, A. M. Siepielski, Disentangling ecologically equivalent from neutral species: The mechanisms of population regulation matter. J Anim Ecol 88, 1755–1765 (2019).

66. A. E. Magurran, B. H. Seghers, A cost of sexual harassment in the guppy, *Poecilia reticulata*. Proc. R. Soc. Lond. B 258, 89–92 (1994).

67. T. A. F. Long, A. Pischedda, A. D. Stewart, W. R. Rice, A cost of sexual attractiveness to high-fitness females. PLoS Biol 7, e1000254 (2009).

68. S. F. Chenoweth, N. C. Appleton, S. L. Allen, H. D. Rundle, Genomic evidence that sexual selection impedes adaptation to a novel environment. Curr Biol 25, 1860–1866 (2015).

69. R. García-Roa, M. Serra, P. Carazo, Ageing via perception costs of reproduction magnifies sexual selection. Proc. R. Soc. B 285, 20182136 (2018).

70. M. J. F. Martins, T. M. Puckett, R. Lockwood, J. P. Swaddle, G. Hunt, High male sexual investment as a driver of extinction in fossil ostracods. Nature 556, 366–369 (2018).

71. M. A. McPeek, L. Shen, J. Z. Torrey, H. Farid, The tempo and mode of three dimensional morphological evolution in male reproductive structures. Am Nat 171, E158–E178 (2008).

72. H. M. Robertson, H. E. Paterson, Mate recognition and mechanical isolation in *Enallagma* damselflies (Odonata: Coenagrionidae). Evolution, 243–250 (1982).

73. D. K. Dowling, B. R. Williams, F. Garciaū Gonzalez, Maternal sexual interactions affect offspring survival and ageing. J Evol Biol 27, 88–97 (2014).

